# Biomass increase under zinc deficiency caused by delay of early flowering in *Arabidopsis*

**DOI:** 10.1101/166884

**Authors:** Xiaochao Chen, Uwe Ludewig

## Abstract

Plants generally produce more biomass when all nutrients are available in sufficient amounts. In addition to environmental constraints, genetic and developmental factors, such as the transition from vegetative to reproductive growth, restrict maximal yield. Here we report the peculiar observation that a subset of early flowering *Arabidopsis thaliana* accessions produced larger shoot rosette diameters when grown in zinc (Zn)-deficient conditions, compared with Zn-sufficient conditions. This was associated with early flowering that restricted the leaf length under Zn sufficiency. Zinc deficiency repressed *FLOWERING LOCUS T (FT)* expression, a major regulator of flowering. Repression or loss of *FT* increased the rosette diameter by a delay of the transition to flowering, a longer phase of leaf proliferation and increased leaf number. The transition to flowering reduced, but not terminated, the proliferation of established leaves. The size of individual leaf mesophyll cells was not affected by Zn deficiency or loss of *FT*, indicating that the larger rosette diameter was caused by maintained proliferation of vegetative tissue. As a consequence, early flowering accessions under Zn deficiency grew larger rosette diameters due to a delay of flowering, which explains the unusual increase of vegetative biomass under nutrient deficiency.

**Highlight:** An increase in biomass of some *Arabidopsis* accessions under Zn-deficiency is caused by retardation of flowering, prolonging vegetative growth.

**Abbreviations:**

DAS: days after sowing
GWA: genome-wide association
SNP: single nucleotide polymorphism
Zn: zinc

## Introduction

Zinc (Zn) is an essential micronutrient for plants and humans. It is a structural component of many catalytic enzymes and transcription factors, so that specific diseases are associated with its deficiency (Broadley *et al*., 2007; Chasapis *et al*., 2012; Marschner, 2011). The Zn bio-availability in many natural soils and crop production systems is low, often as a result of high CaCO_3_ content and alkaline soil pH. As a consequence, many food products are low in Zn, causing malnutrition in humans (Cakmak, 2008).

In *Arabidopsis*, over 2000 Zn-related genes are annotated, primarily with catalytic and transcriptional regulator activities (Broadley *et al*., 2007). Zn-responsive key genes and transporters involved in Zn uptake and translocation have been reported using molecular and genetic tools (Sinclair and Kramer, 2012). Severe Zn deficiency leads to reduced, abnormal leaf and seed growth, in addition to impaired flower development (Talukdar and Aarts, 2007). Generally, plant development and flowering can be delayed when a certain nutrient is unavailable to the plant, but general nutrient deficiency in *Arabidopsis* has been associated with acceleration of flowering (Kolar and Senkova, 2008). In the ornamental plant *Pharbitis nil*, poor nutrition promotes flowering and this is correlated with elevated expression of the major flowering integrator *FLOWERING LOCUS T (FT)* (Wada *et al*., 2010).

The appropriate decision for flowering in annual plants is crucial for their lifespan and is under very complex genetic control, with over 360 genes implicated (Andres and Coupland, 2012; Bouche *et al*., 2016). The transition to flowering depends on several endogenous and environmental signals, which are best studied in *Arabidopsis thaliana*. Several flowering pathways have been identified, such as photoperiod, temperature, vernalization, gibberellin and sugar pathways (Andres and Coupland, 2012; Bouche *et al*., 2016; Capovilla *et al*., 2015). Flowering signals converge in the activation of the *FT* gene in source leaves. Its translated gene product (florigen) is phloem mobile and is translocated to the shoot apical meristem, where *FT* dimerizes with the transcription factor *FLOWERING LOCUS D (FD)* to activate another central integrator, *SUPPRESSOR OF OVEREXPRESSION OF CONSTANS 1 (SOC1)*. This terminates the vegetative fate of the apical meristem and initiates flower development (Corbesier *et al*., 2007; Jaeger and Wigge, 2007; Notaguchi *et al*., 2008). While the transition to the flowering fate in the apical meristem is well explained by *FT* (Amasino, 2010; Andres and Coupland, 2012), its role in the regulation and termination of vegetative leaf growth is less well understood (Melzer *et al*., 2008; Shalit *et al*., 2009). In already established leaves, the final leaf size and the rosette diameter are ultimately controlled by the complex coordination of primordium size, cell proliferation and cell expansion (Gonzalez *et al*., 2012; Powell and Lenhard, 2012). The maintenance of vital meristematic regions in leaves is essential to obtain maximal leaf growth and size (Gonzalez *et al*., 2012; Powell and Lenhard, 2012).

In preliminary experiments we initially observed that in a Zn-deficient soil-sand mixture, some *Arabidopsis* plants finally produced more shoot biomass than in a Zn-sufficient soil. To uncover the genetic basis for this unusual behavior, we grew 168 *Arabidopsis* accessions in low Zn and Zn-amended soil-sand mixtures and quantified the rosette diameters of these accessions as a proxy for shoot size and leaf growth (length). Unexpectedly, Zn deficiency prolonged vegetative growth in some early-flowering accessions, leading to larger plants, an effect potentially genetically associated with *FT*. Loss-of-function mutants of flowering genes also differed in their rosette size in a Zn-dependent manner. While it is well accepted that the transition to flowering in the shoot apical meristems ultimately terminates further vegetative growth, our results uncover that FT only gradually represses already established vegetative leaves, at least in a subpopulation of *Arabidopsis*.

## Results

### Natural variation of rosette diameter and its response to zinc deficiency

To explore the natural variation and differential plant growth as the function of differential Zn availability in soil, 168 *Arabidopsis thaliana* accessions were grown in a fertilized soil-sand mix without (-Zn) or with added ZnSO_4_ (+Zn) in the greenhouse. The set of accessions included six main populations (Central Europe, Northern Europe, Iberian Peninsula, Mediterranean, Central Asia and North America) and three small populations (Cape Verde, Canary Islands and Japan) (Supplemental Figure S1, Supplemental Table 1) (Chen *et al*., 2016; Stetter *et al*., 2015).

There was substantial natural variation in the vegetative shoot growth and rosette diameter among these accessions in +Zn and -Zn, which represented Zn sufficiency and mild Zn deficiency conditions, respectively (Figure 1A). The rosette diameter, a proxy for the maximal vegetative leaf length (with petiole), ranged from 1.3 cm to 12.6 cm in +Zn, with a major distribution peak of around 10 cm. In -Zn, the rosette diameter ranged from 2.3 cm to 9.6 cm, with the majority of accessions having a rosette of around 7 cm (Figure 1A; Supplemental Table S2). One-way ANOVA indicated that significant genetic differences were observed among accessions (p<2e-16), in both conditions (Supplemental Table S3). Broad-sense heritability of rosette diameter was 0.69 for +Zn, but only 0.38 for -Zn (Supplemental Table S3). As expected, the rosette diameter in -Zn highly correlated with that in +Zn (p<2.5e-8, Supplemental Figure S2), as the leaf size is principally genetically controlled. Interestingly, the data suggest that the 168 accessions adjust their growth differently depending on Zn availability. Most accessions produced smaller final rosette diameters under -Zn, as expected when an essential nutrient is limiting for leaf growth. However, the others ended up with larger rosette diameters under -Zn (Figure 1A-B). These accessions included the widely used accession Col-0.

**Figure 1:**
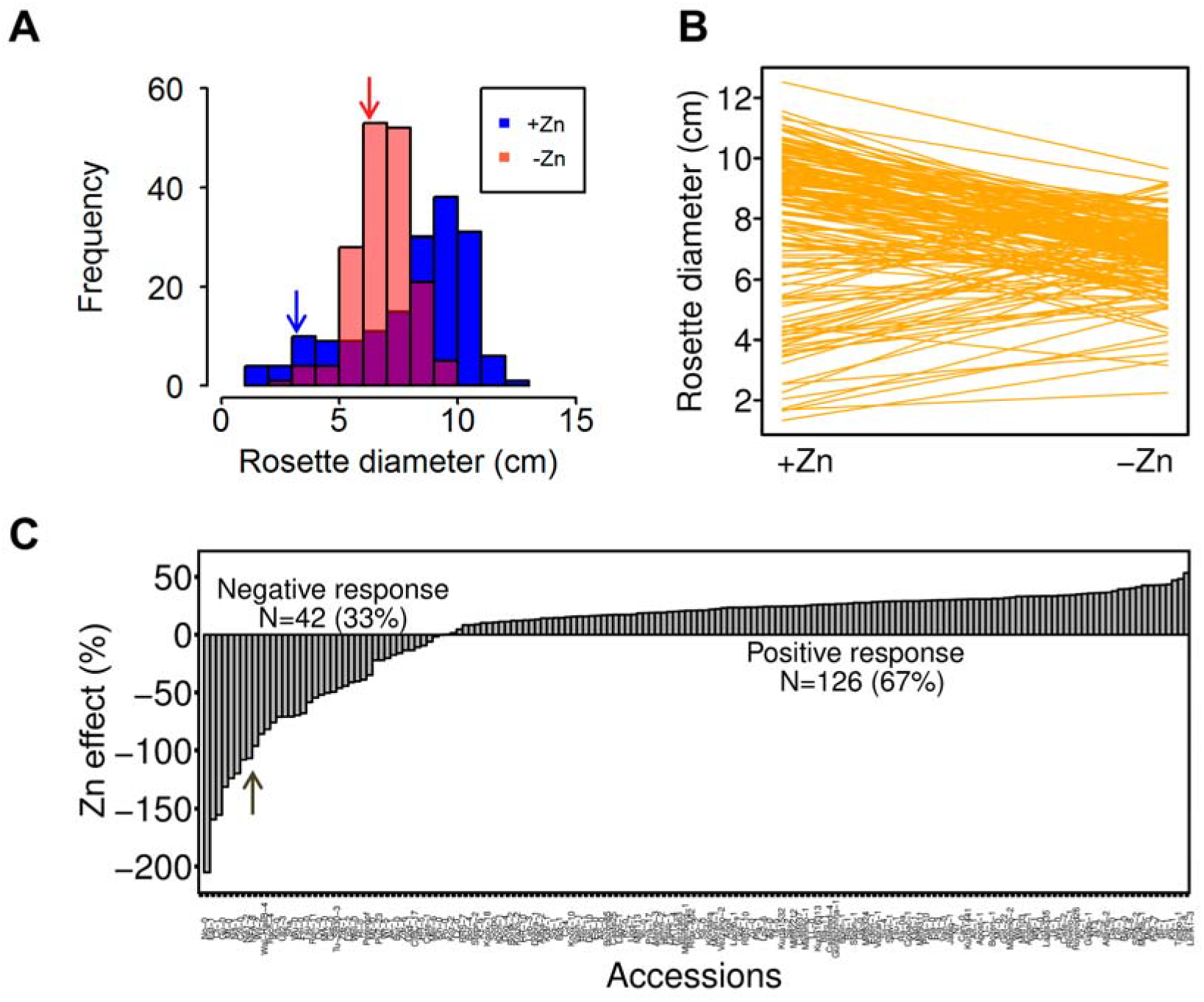
Natural variation of rosette diameter and its response to Zn deficiency. **A**, Distribution of rosette diameter for 168 accessions grown under control (+Zn) and Zn deficiency (-Zn) conditions. Data plotted are mean + SD, n=6. **B**, Reaction norms of rosette diameter in +Zn and -Zn. **C**, Natural variation of Zn effect. 168 accessions were divided into negative response and positive response. Arrows in **A** and **C** indicate the accession Col-0.

To quantify this interesting phenomenon, we defined the relative reduction of the rosette diameter due to Zn deficiency as ([(+Zn) – (-Zn)] * 100 / +Zn) and called this number “Zn effect” on the final rosette diameter. This “Zn effect” is positive for plants with increased rosette diameter in Zn-amended soil. A positive “Zn effect” thus represents the situation in which higher availability of an essential nutrient increases the final plant size, likely because this element was growth limiting. However, the Zn effect is negative for plants with larger rosette diameter in Zn-deficient soil. This negative “Zn effect” thus represents a situation in which other factors than missing Zn limited further growth. 126 accessions (67% of 168 accessions, e.g. Sf-2) had decreased rosette diameter in Zn-deficient soil (Figure 1C, positive Zn effect on leaf size). By contrast, the other 42 accessions (33% of 168 accessions, e.g. Col-0) had a decreased rosette diameter, despite having sufficient Zn in soil (Figure 1C, negative Zn effect). The average Zn effect was -64.5% for the negative responses and 24.8% for the positive responses. The Zn effect was determined by +Zn soil (r=0.85, p<2.2e-16), rather than -Zn soil (r=0.06, p=0.4478, Supplemental Figure S2), indicating that the Zn effect was not primarily due to different growth and development under Zn deficiency. Rather, strong heterogeneity of growth and flowering time in the population was observed in +Zn and this heterogeneity was lost in -Zn.

### Genome-wide association mapping for the Zn effect

To identify the underlying genetics of the Zn effect on rosette diameter, we carried out genome-wide association (GWA) mapping with 162 accessions for which high density single nucleotide polymorphisms (SNPs) were available. A multi-locus mixed model was implemented to eliminate noise of population structure (Segura *et al*., 2012). A stringent p-value cutoff with 5% false discovery rate (FDR) was set to quantify the significance.

The GWA did not identify any significant SNP for the rosette diameter under both +Zn and –Zn conditions (data not shown). However, four significant candidate SNPs were observed for the Zn effect: *Chr1_16581335, Chr1_23946852, Chr1_24341704* and *Chr4_11742318* (Figure 2; Supplemental Figure S3 & S4). Although none of these loci qualified for a distinguished high quality candidate locus with numerous co-segregated SNPs, we noted that the 48 transcripts located +/- 20 kb of these four SNPs were enriched in genes annotated as being involved in flowering or were overrepresented in being expressed during reproduction (Supplemental Table 4). *AT1G43800* (*FTM1, FLORAL TRANSITION AT THE MERISTEM1*) and *AT1G65480 (FT, FLOWERING LOCUS T)* locate 535 bp and 7770 bp distant of the significant SNPs *Chr1_16581335* and *Chr1_24341704*, respectively. *FTM1* is activated independently of *FLOWERING LOCUS T (FT)* and *SUPPRESSOR OF OVEREXPRESSION OF CONSTANS1 (SOC1)* during the floral induction (Torti *et al*., 2012). *FT* is central to flowering and the allele G at *Chr1_24341704* in the vicinity of *FT* was associated with the negative Zn effect (Supplemental Figure S4). It was less frequently represented in the population than the allele A.

**Figure 2:**
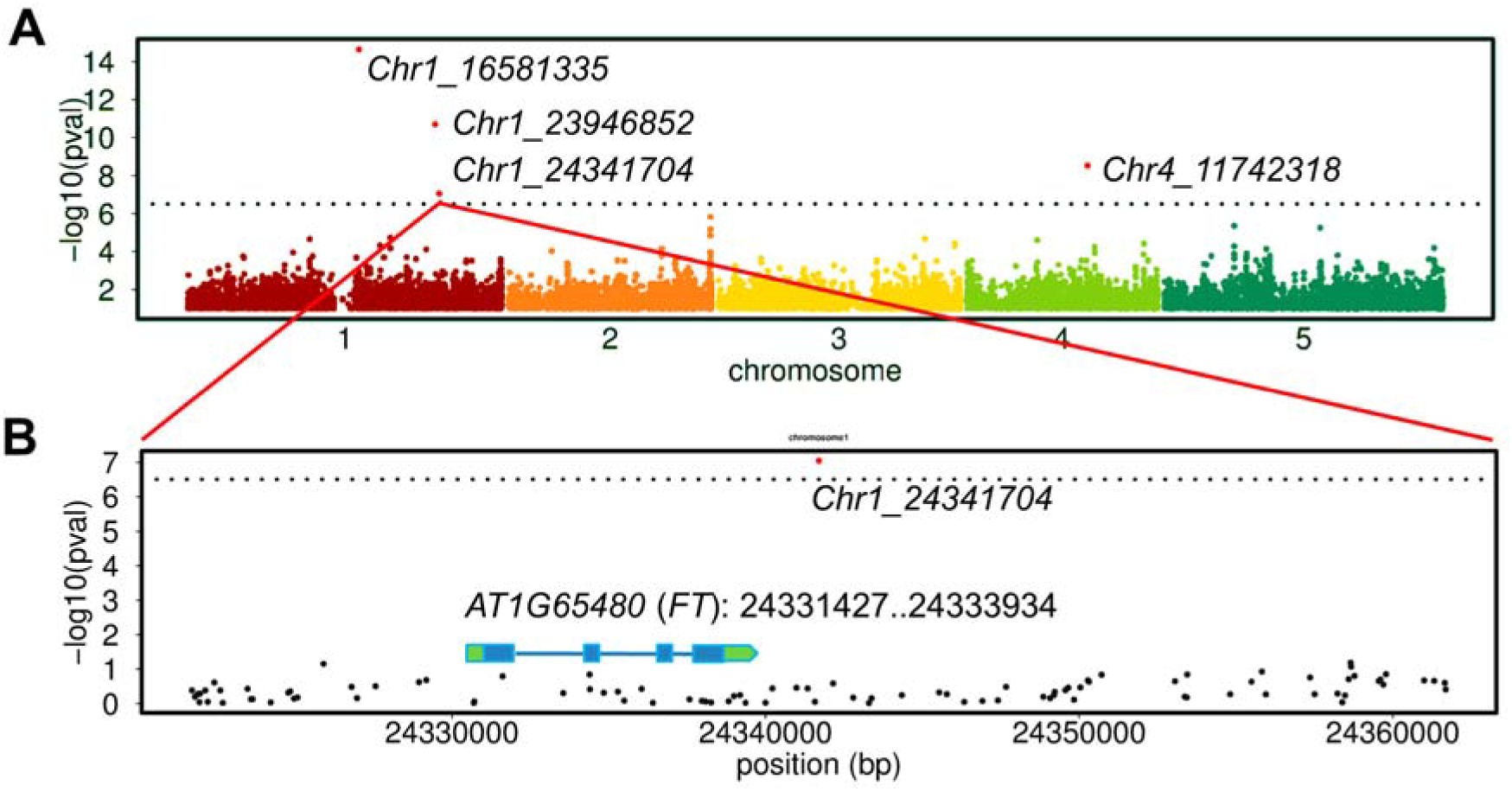
Genome-wide association mapping of Zn effect. **A**, Manhattan plot for Zn effect of rosette diameter. The 5% FDR threshold was denoted by a dashed line. Red dots indicate significant SNPs. **B**, +/- 20 kb windows fine map of significant SNP *Chr1_24341704*, which located around 7.8 kb distance of *FT*.

### Relationship of rosette diameter with flowering and Zn

As flowering genes were identified as potential candidates in the GWA, we assumed that flowering time is potentially correlated with the final rosette diameter, two traits with typically minor correlation (Atwell et al., 2010). Indeed, flowering time was highly correlated with the rosette diameter in negative-response accessions, which flowered typically earlier (Figure 3A). However, no significant correlation with flowering was found in the accessions with positive Zn effect. Moreover, we examined how the central flowering integrator *FT* was affected by Zn via measuring its expression level, including four negative-response accessions (Col-0, Po-0, Ct-1, No-0) and four positive-response accessions (Lerik1-3, Koz-2, Sf-2, Cvi-0). RNA was extracted at 14 DAS (days after sowing), when the plants were still in vegetative growth. -Zn greatly reduced *FT* expression in all accessions (Figure 3B). In addition, negative-response accessions presented higher *FT* level compared to positive-response accessions, in agreement with early flowering and *FT* involved in the Zn effect. The same gene expression pattern was observed for another central floral integrator, *SOC1* (Supplemental Figure S5).

**Figure 3:**
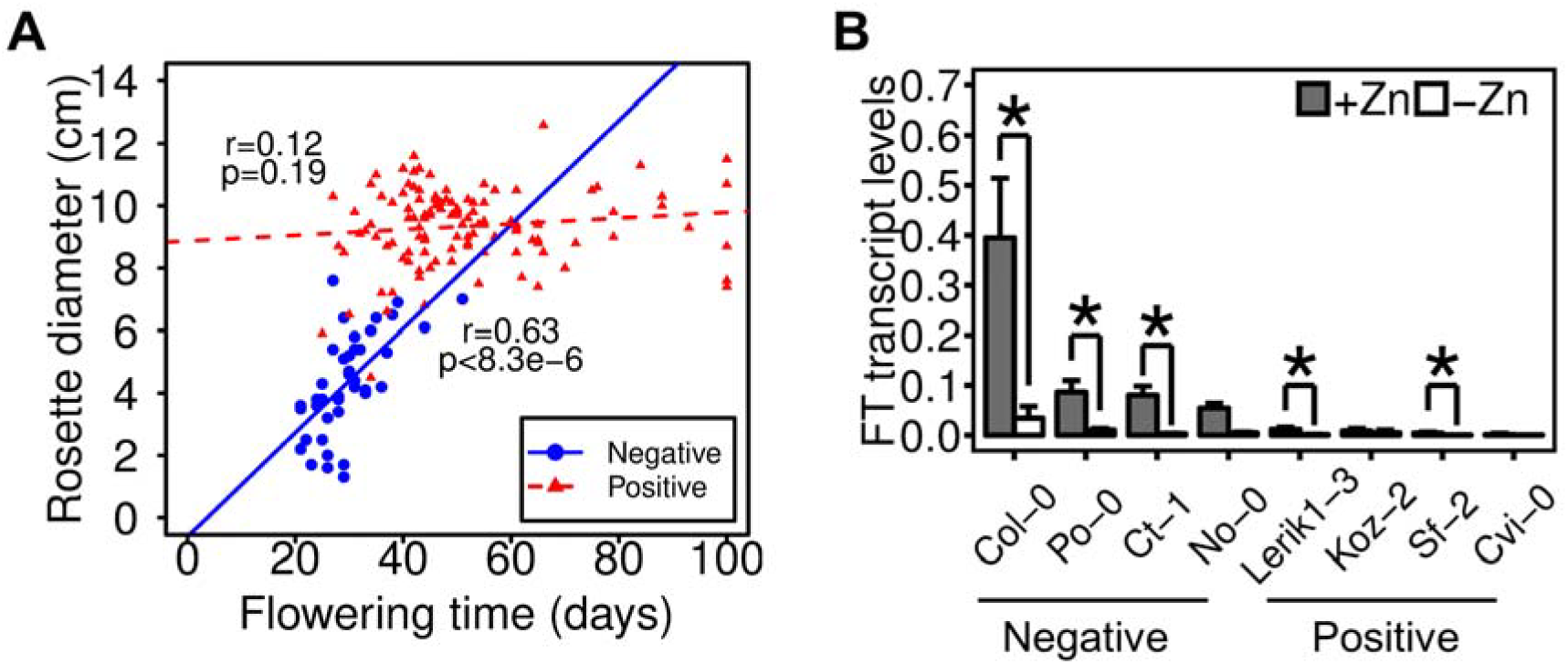
Relationship of rosette diameter with flowering time and Zn. **A**, Correlation between flowering time (+Zn) and rosette diameter in negative-response accessions and positive-response accessions. **B**, *FT* transcript levels in negative-response (Col-0, Po-0, Ct-1, No-1) and positive-response accessions (Lerik1-3, Koz-2, Sf-2, Cvi-0). Gene expression was referenced to *SAND* and *PDF2* genes. * denotes the significant difference at p<0.05 level.

### Genetics of early flowering, rosette size and relation with Zn

The phenotypic differences within a batch of Col-0 plants are shown in Figure 4A; obviously, -Zn repressed and delayed flowering. Further genetic analysis concentrated on this negative-response accession (Zn effect was -119 %), with strong *FT* and *SOC1* inhibition by –Zn (Fig. 4B). To get more insight whether photoperiod, sugar, vernalization, gibberellin or ambient temperature pathways were affected by Zn availability (Andres and Coupland, 2012), we checked the expression of key genes representative of these flowering pathways during vegetative growth. The expression of the diurnal integrator *CONSTANS (CO)* and the trehalose biosynthesis enzyme *TREHALOSE-6-PHOSPHATASE SYNTHASE 1* (*TPS1*) was not different. Furthermore, the transcriptional repressors *FLOWERING LOCUS C (FLC), FLOWERING LOCUS M (FLM)* and *SHORT VEGETATIVE PHASE (SVP)*, the latter two being inhibitors of the elevated temperature-induced early flowering pathway, were all unaffected by Zn. Finally, the flowering promoting *GIBBERELLIN 3-OXIDASE1* (*GA4*), as well as the expression of *FTM1*, identified from the GWA, were also not different between -Zn and +Zn (Fig. 4B).

**Figure 4:**
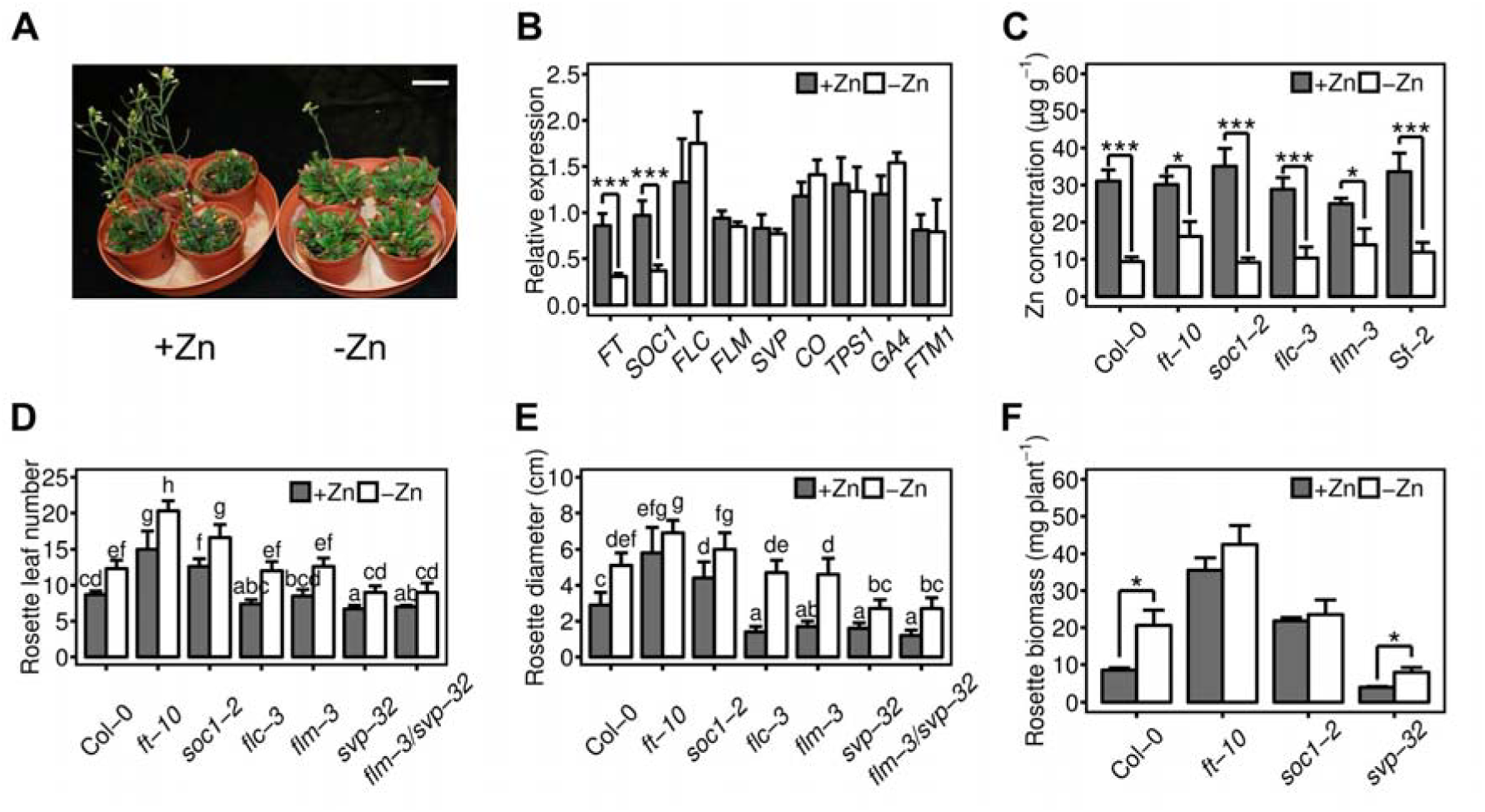
Genetic basis of Zn regulation of flowering time and rosette diameter. **A**, Col-0 plants grown in +Zn and -Zn soils. Scale bar: 5cm. Note that -Zn delayed flowering. **B**, Relative expression level for central flowering genes in Col-0 plants at vegetative growth stage (14 DAS). Replicated data were normalized to reference genes *SAND* and *PDF2*. **C**, Shoot Zn concentrations in –Zn and +Zn conditions. **D**, Rosette leaf number and **E**, rosette diameter, at bolting stage in wild-type (Col-0) and flowering null mutants in Col-0 background. **F**, vegetative biomass in Col-0 and mutants, *** denote p<0.001. Different small letters between columns denote significant difference at p<0.05 level. Data plotted are mean + SD.

Next we tested how mutant null alleles of key flowering genes in the Col-0 background responded to different Zn availability, including *ft-10, soc-1-2* and *flc-3*. Since early flowering appeared to be sensitive to Zn, we also included mutants in the elevated temperature-induced early flowering pathway, *flm-3 and svp-32* (Balasubramanian *et al*., 2006; Pose *et al*., 2013). The shoot Zn concentrations for all these mutants, its corresponding wild type Col-0 and the accession Sf-2 were below 20 μg/g, a typical deficiency threshold in many plant species. Shoot Zn massively differed between -Zn and +Zn conditions (Figure 4C). Furthermore, typical Zn-deficiency responsive genes, such as the Zn transporter genes *ZIP1, ZIP4, ZIP5* and *IRT3*, were induced under -Zn conditions in both Col-0 and Sf-2, indicating the activation of known physiological Zn-deficiency responses (Supplemental Figure S6). The flowering time was quantified at bolting stage by counting rosette leaf numbers rather than growth days, to eliminate the influence of growth speed (Balasubramanian *et al*., 2006; Pose *et al*., 2013). In agreement with previous studies, *ft-10* and *soc-1* had delayed flowering time, whereas *svp-32* and *flm-3*/*svp-32* accelerated flowering (Figure 4D). Meanwhile, *flc-3* and *flm-3* had similar flowering time as Col-0, which is likely explained by the weak *FLC* and *FLM* alleles in the accession Col-0 (Balasubramanian *et al*., 2006; Lempe *et al*., 2005). Nevertheless, Zn deficiency delayed flowering in all genotypes by a few days, irrespective of their widely different overall flowering time (Figure 4D). Interestingly, synchronously with this delay of flowering in –Zn, the rosette diameter of all genotypes increased significantly, except for *ft-10*, where only a minor trend, but no significant increase in rosette diameter was observed (Figure 4E). The larger rosette diameter of Col-0 in – Zn, as well as the increased number of leaves due to longer vegetative growth finally translated into larger total vegetative biomass, as compared with the Zn-amended control condition. The same was true for *svp-32*, while the final vegetative biomass of *ft-10* and *soc-1* did not differ between –Zn and +Zn (Figure 4F).

### Manipulation of the zinc effect by growth temperature or vernalization in two genetic backgrounds

To further get mechanistic insight into the Zn effect, we next asked whether the Zn effect could be eliminated or induced by environmental manipulation of the flowering time. Therefore, the early flowering accession Col-0 and the late-flowering accession Sf-2 were either grown at low temperature (16°C, Col-0) or were vernalization pre-treated before growth at 23°C, to promote flowering (Sf-2), respectively. Because of the causal roles of *FT* and *SOC1* genes in regulating flowering, their expression in the various conditions was monitored. Expression was strongly reduced by Zn deficiency after 2 weeks, irrespective of the growth temperature in Col-0 (Figure 5 A-B). However, this difference in gene expression did not remain after 7 weeks at 16°C, when plants still were in vegetative stage (Figure 5 C). Similarly, in Sf-2, the reduced expression of *FT* and *SOC1* in –Zn was stronger at 2 weeks, but only a minor *FT* expression difference was apparent after 7 weeks of vegetative growth (Figure 5 D-F). In agreement with this gene expression pattern, Col-0 flowered after 3 weeks at 23°C, but flowering time was delayed to over 7 weeks at 16°C. Interestingly, the delay of flowering in -Zn was entirely lost at 16°C, leading to equal rosette diameters in –Zn and +Zn conditions (Figure 5G-H). Furthermore, the rosette diameter of Col-0 plants grown in +Zn was larger at 16°C compared to 23°C, because flowering terminated the vegetative growth in plants grown at the higher temperature. On the other hand, vernalization of Sf-2 was able to accelerate flowering time to 3 weeks, although without vernalization, this accession flowered only later than 7 weeks. Accordingly, the flowering time and rosette diameter of non-vernalized Sf-2 were indistinguishable between different Zn supplies (Figure 5G-H). By contrast, the early flowering of vernalized Sf-2 was delayed in –Zn and, consequently, the rosette diameter was slightly increased in –Zn. Although all Sf-2 plants (vernalized and non-vernalized) were grown at 23°C, the vernalized plants finally had smaller rosette diameters, because they flowered earlier (Figure 5G-H).

**Figure 5:**
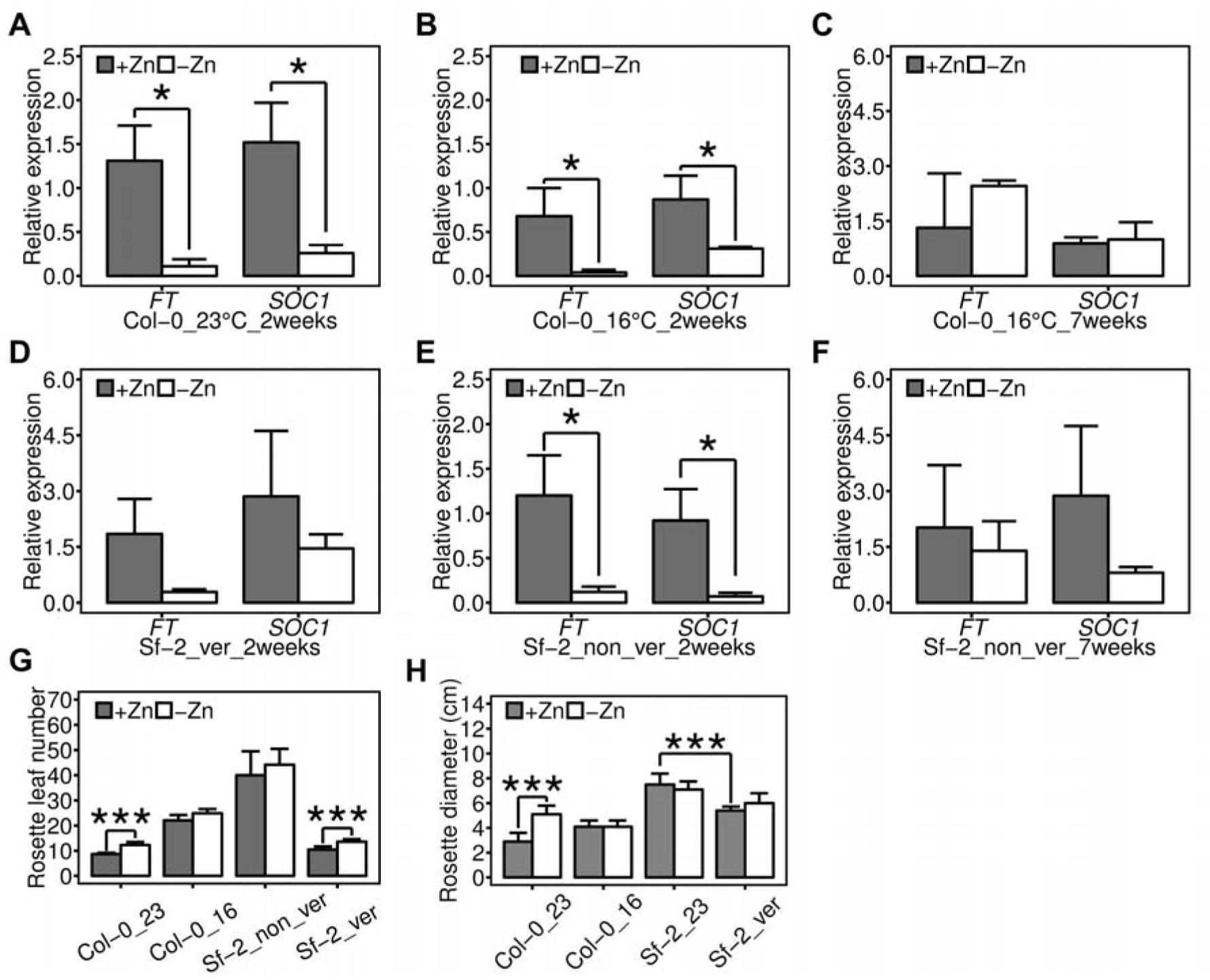
Low temperature and vernalization affect the Zn-dependence of flowering in natural accessions. **A-C**, Relative *FT* and *SOC1* expression levels of Col-0 grown at 23 and 16 °C. **D-F**, Relative *FT* and *SOC1* expression levels of Sf-2 grown with and without vernalization process. Vernalization indicates the process of 3-weeks cold treatment before normal growth. **G**, Rosette leaf number and **H**, rosette diameter of Col-0 and Sf-2 at bolting stage. “ver” means vernalization. * and *** denote p<0.05 and p<0.001, respectively. Data plotted are mean + SD.

### Leaf cell proliferation contributed to the differences in leaf length

The final leaf length (and rosette diameter) are ultimately controlled by the complex coordination of leaf primordium size, differential cell proliferation in different areas of the leaf and cell expansion (Gonzalez *et al*., 2012; Powell and Lenhard, 2012). To get insight into the underlying mechanism behind the increased final leaf length in -Zn (Figure 6A), the leaf mesophyll cell size and cell numbers were microscopically quantified at two stages, 19 DAS and 33 DAS, using confocal imaging (Figure 6B). Plants in –Zn and +Zn conditions did not flower at 19 DAS, but fully flowered at 33 DAS. The late flowering *ft-10* mutant was also analysed. Interestingly, the cell size was not distinguishable at 19 DAS for Col-0 or at 33 DAS in Col-0 and *ft-10*, irrespective whether grown in -Zn or +Zn (Figure 6C). Since Col-0 leaves differed in size at 33 DAS depending on Zn (and differed from *ft-10*, Figure 6A), this indicated that the cell number differences (and thus cell proliferation in established leaves) mainly contributed to the differences in the leaf length.

**Figure 6:**
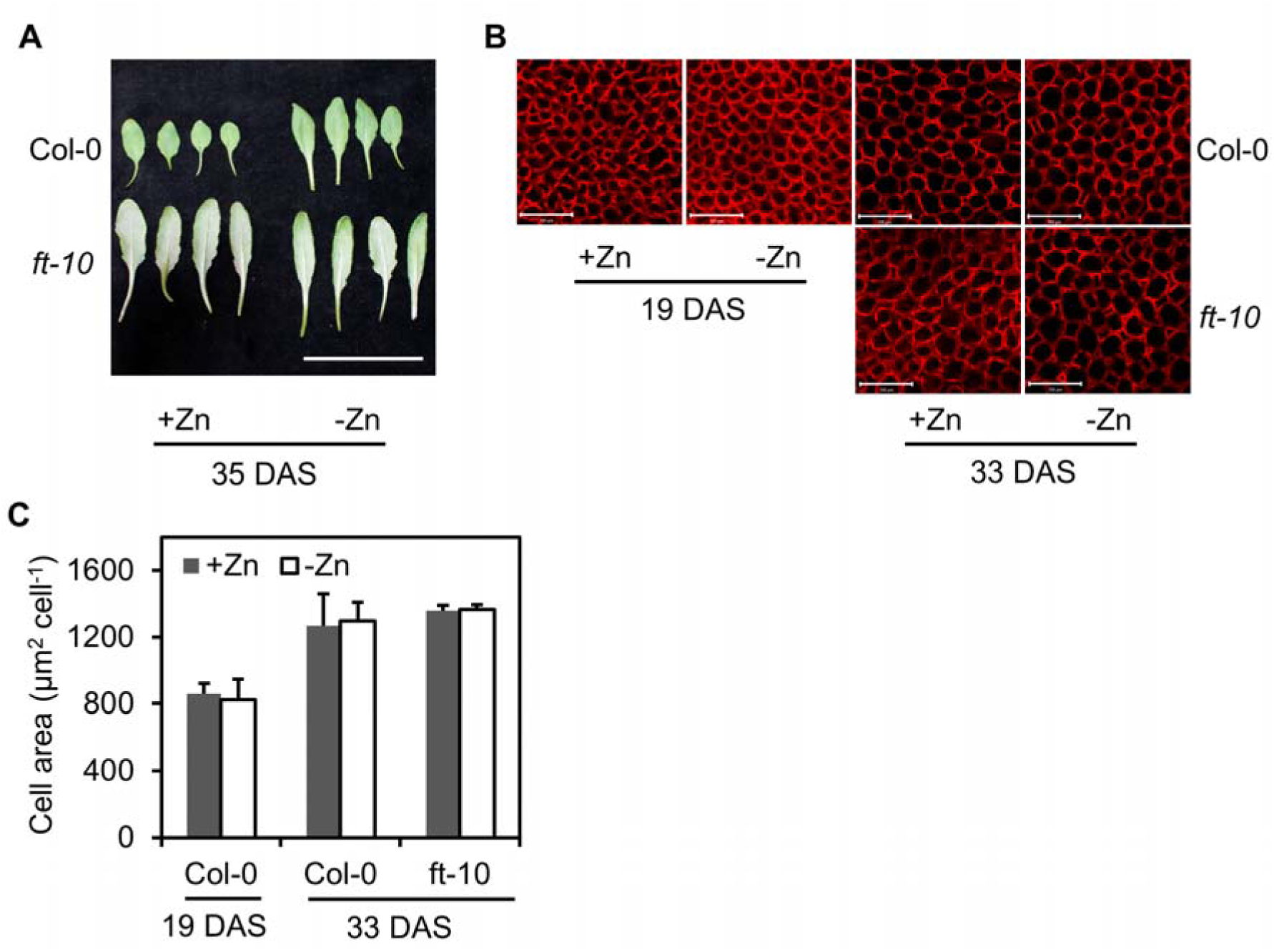
Quantification of cell number in leaves. **A**, First four leaves at 35 DAS (days after sowing). Scale bar: 5cm. **B**, Confocal mesophyll cell images after staining with propidium iodide of wild type and *ft-10*. Scale bar: 100 μm. **C**, Average cell size of first four wild type and *ft-10* leaves at 19 DAS and 33 DAS. Note that plants have not flowered at 19 DAS and thus used as the background, and plants finished flowering at 33 DAS. Data plotted are mean + SD.

### Gradual repression of leaf growth rate via zinc-promoted flowering

The transition of vegetative to reproductive growth by the induction of flowering is commonly centred to the apical shoot meristem, but how the growth of already established leaves is also finally terminated and ends in senescence, is little investigated. In order to quantify the onset of leaf growth rate inhibition by Zn and FT, we documented the growth curve of the rosette diameter, as well as the petiole length in Col-0 and *ft-10* plants. The rosette diameters were initially indistinguishable between +Zn and -Zn, in agreement with the assumption that the essential Zn was not growth limiting at this stage. However, the rosette diameter differed just after the transition to flowering in the apical meristem had occurred (Figure 7A), consistent with an immediate repressing signal transmitted at the time of flower initiation in +Zn (21 DAS, Figure 7A). Interestingly, the longitudinal leaf growth rate was not terminated, but only reduced, after the transition to flowering. The longitudinal leaf growth rate in -Zn remained larger than in +Zn, until termination at around 33 DAS (Figure 7A). Such a repression of leaf growth was lost in *ft-10*, which did not flower before 32 DAS, leading to strong leaf expansion irrespective of Zn supply (Figure 7B). The petiole elongation showed a similar pattern as the entire rosette diameter in the response to Zn deficiency, suggesting a similar restriction mechanism in the petiole and in the leaf blade (Figure 7 A-B).

**Figure 7:**
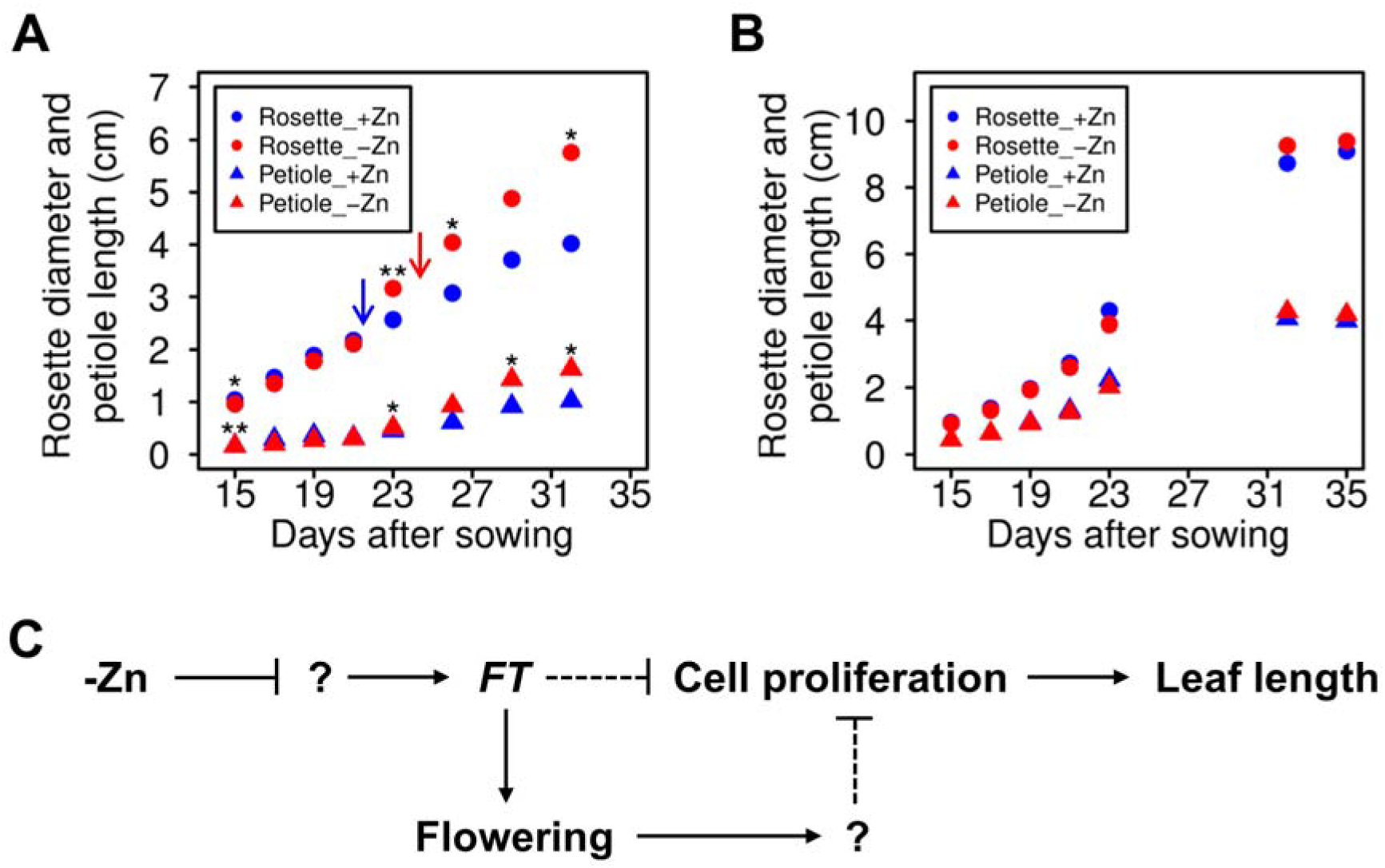
Zn regulation of flowering time and leaf length. **A**, Growth curve of rosette diameter (circle) and petiole size (triangle) in Col-0 under +Zn (blue) and -Zn (red) conditions. * and ** denote p<0.05 and p<0.01. Arrows indicate the flowering time in +Zn (blue) and -Zn (red). **B**, Growth curve of *ft-10*. **C**, Working model of Zn regulation of flowering time and leaf length in early-flowering *Arabidopsis*. Arrows and block lines denote activation and repression, respectively. Dashed lines indicate putative regulation.

## Discussion

Substantial natural variation has been previously reported in *Arabidopsis thaliana*, including rosette diameter, leaf shape, ion content, flowering time and many other physiological characteristics (Salt *et al*., 2008; Weigel, 2012). However, only limited knowledge has been obtained to understand how the natural variation growth phenotypes are influenced by nutrient deficiency. Here, we investigated the rosette diameter of 168 *Arabidopsis* accessions grown in +Zn (control) and –Zn (Zn deficiency) soil-sand mixes. Randomly chosen plants grown on the Zn-deficient soil-sand mix had only Zn concentrations of ~10-13 μg/g in their shoot tissue, but did not show visual signs of Zn deficiency. As expected, the rosette diameter was highly variable among these 168 accessions in both conditions (Figure 1). Surprisingly, not all accessions reduced their rosette diameters in –Zn, which was inconsistent with our naive expectation that plants must grow better and produce more biomass when all essential nutrients are adequately supplied (Marschner, 2011). To mechanistically unravel this interesting finding, we quantified the “Zn effect” by calculating the relative difference in rosette diameter, to indicate how different accessions respond to Zn deficiency. The 168 accessions were divided into 126 positive Zn effect accessions (e.g. Sf-2) and 42 negative accessions (e.g. Col-0). In the positive Zn effect accessions the addition of Zn increased the plant diameter, which is expected if an essential nutrient is growth limiting. However, larger rosette diameters in –Zn were found in the negative Zn effect accessions, compared to +Zn conditions, suggesting that other factors apart from missing Zn reduced the growth.

We performed multi-locus mixed model genome-wide association (GWA) mapping for the Zn effect (Figure 2). Although four significant SNPs were associated with the Zn effect, of which two were close to flowering genes *FT* and *FTM1*, none of these loci qualified for a convincing, strong hit with multiple linked co-segregated significant SNPs in the vicinity ~20 kb, which are expected for causal loci. However, any significant SNP hit may deserve further analysis, although a larger population is likely required to further validate the candidate SNPs identified. The suggested link to flowering potentially indicated that we unintentionally screened for flowering differences in the population, as flowering was indeed affected by Zn. In earlier population studies with *Arabidopsis*, the flowering time and rosette diameter had only minor correlation (Atwell *et al*., 2010). Here, surprisingly, we found a strong positive correlation between flowering time and rosette diameter, but only in the 42 negative-response, early flowering accessions (Figure 3). No significant correlation was found in the 126 positive-response accessions. Under the conditions used, early flowering was prominent in +Zn, but lost in –Zn. Late flowering accessions probably maintained a longer vegetative growth phase and leaves were already fully expanded before flowering, but it is worth to note that some among the negative Zn effect accessions also flowered very early. This potential connection of Zn with flowering was analysed in more detail. *FTM1* expression was unchanged by Zn supply, but the *FT* expression was generally repressed under Zn deficiency (Figure 3 & 4).

In several key mutants in the Col-0 background that are affected in (early) flowering, Zn consistently delayed flowering, in agreement with the observations in natural accessions with different flowering time. This indicated a strong genetic connection with flowering time, but failed to identify all convincing causal targets of the Zn effect. General poor nutrient supply had been shown to accelerate flowering in *Arabidopsis* (Kolar and Senkova, 2008). The macronutrients nitrate and phosphate act in an antagonistic way, as nitrate deficiency promotes and low phosphate delays flowering (Kant *et al*., 2011). At higher concentrations, nitrate promotes flowering independent of the phytohormone gibberellin, which integrates several environmental stimuli and acts downstream of other known floral induction pathways (Castro Marin *et al*., 2011). Thus, the effects of nutrient deficiencies on flowering are not uniform. Flowering is also promoted by mineral stress (50 μM cadmium, toxic for *Arabidopsis*) via up-regulation of *CONSTANS* (*CO*) and *FT* (Wang *et al*., 2012). Zn deficiency generally repressed *FT* expression, even in late flowering accessions and delayed flowering in some genotypes. Interestingly, Zn deficiency consistently led to larger rosette diameters in negative Zn effect accessions. Because of the prolonged vegetative growth time, longer and more leaves were established, so that –Zn grown plants finally accumulated more vegetative biomass then +Zn-grown plants (Figure 4). The canonical flowering pathways, such as photoperiod, temperature, vernalization, sugar and gibberellins, were apparently unaffected by Zn (Figure 4). The reduced expression of *SOC1* in –Zn could be just a consequence of *FT* repression, as *SOC1* is a downstream gene of *FT* (Andres and Coupland, 2012). Plants lacking *FT* flowered later than *soc-1-2* and produced larger rosettes, confirming that *FT* is the central floral integrator. Consequently, Zn also regulated flowering time and rosette diameter in *soc1-2* plants. The direct target of the Zn effect, however, remains unclear, as flowering was even delayed in the very late flowering *ft-10* in –Zn. Some further regulator of *FT* expression might be Zn-dependent. It is noted, however, that the small difference between *ft-10* large rosette diameters in +Zn and –Zn was not significant, suggesting that the Zn effect was largely lost in this mutant and that a SNP close to the *FT* gene was genetically associated with the Zn effect.

While our data suggest that *FT* expression can be delayed in -Zn, there is likely also a “memory” effect induced under different Zn supply, as in individual, genetically identical Sf-2 plants grown under identical conditions, the Zn effect could be introduced by vernalization pre-treatment (Figure 5). Indeed, *FT* expression is epigenetically controlled (Andres and Coupland, 2012). In late-flowering plants, many other factors apart from FT regulate flowering time and restrict organ size, including genes involved in auxin, cytokinin and gibberellin signaling (Bogre *et al*., 2008; Powell and Lenhard, 2012). The target of FT to terminally restrict vegetative leaf growth is, however, unlikely the shoot apical meristem alone, as suggested by the leaf cell numbers and rosette growth curves of Col-0 and *ft-10* plants (Figure 6 & 7). The rosette diameters were initially indistinguishable between +Zn and –Zn (Figure 7), but differed after flowering, indicating that a repressing signal was received in already fully established, 1 cm long rosette leaves in +Zn (21 DAS). This repressing signal could be *FT* itself or a downstream signal, but importantly, the inhibition of cell proliferation that results in less leaf cell numbers in +Zn, was entirely lost in *ft-10*, which is known to have a much larger rosette diameter (Figure 6).

A schematic working model summarizes the dual effects of *FT* in flowering and leaf length restriction (Figure 7C). -Zn repressed *FT* expression in certain early flowering genotypes via an unknown and potentially indirect process, leading to delayed flowering compared to +Zn plants. As a consequence, plants in Zn deficiency produced more and longer leaves via prolongation of vegetative growth. This Zn effect, however, is only found in a subset of the *Arabidopsis* population, namely in early flowering genotypes, and is clearly different from general nutrient deficiencies, which promote flowering. From an ecological perspective, the two strategies to respond to limited Zn may be beneficial in different environments. Restricting further vegetative growth after abiotic stress-induced early flowering, as observed in the negatively Zn-responsive genotypes, will lead to early, but few seeds, while the delay of flowering under –Zn will prolong the vegetation period, increase accumulation of nutrients, that finally are transferred to many, but late seeds. Whether this nutritional regulation of flowering and the concomitant restriction of vegetative growth are relevant in ecosystems and crops, are interesting questions for future research.

## Methods

### Plant material, soil-sand preparation and growth conditions

168 *Arabidopsis thaliana* accessions used in this study are listed in the Supplemental Table S1. Seeds for all accessions were obtained from Dr. Karl Schmid (Germany). The *ft-10, soc1-2, flc-3, flm-3, svp-32, flm-3/svp-32* mutants in the Col-0 background were gifted by Dr. Markus Schmid (Umea, Sweden). All accessions and mutants have been previously described (Balasubramanian *et al*., 2006; Pose *et al*., 2013; Stetter *et al*., 2015).

Soil-sand mixtures of a Zn-scarce soil from a C-horizon of a loess soil (0.7 mg kg^-1^ Zn, pH 7.2) was mixed at 1:1 ratio with quartz sand (0.6-1.2 mm diameter), which was washed with HCl (rinsed with tap water, pH<1 adjusted with HCl, incubated for one day, rinsed with deionized water to pH>5) to wash out trace nutrients, biological contaminations and dust. The soil-sand mix was fertilized with 1.1 g kg^-1^ NH_4_NO_3_, 0.9 g kg^-1^ K2SO_4_, 2.1 g kg^-1^ MgSO_4_ and 1.6 g kg-1 Ca(H_2_PO_4_)_2_. 200 g of soil-sand per plant (or 120 g for qRT-PCR experiments and mutant experiments) was placed in the pots before watering with 7-8 ml micronutrients, according to a modified Hoagland’s solution (1 mM NH_4_NO_3_, 1 mM KH_2_PO_4_, 0.5 mM MgSO_4_, 1 mM CaCl_2_, 0.1 mM Na_2_EDTA-Fe, 2 μM ZnSO_4_, 9 μM MnSO_4_, 0.32 μM CuSO_4_, 46 μM H_3_BO_3_, 0.016 μM Na_2_MoO_4_). In addition, 3 mg kg^-1^ ZnSO_4_ was added into soil for the control treatment (+Zn).

Seeds were stratified at 4°C for 3 days to promote germination. Plants were cultivated in greenhouse (GWA, during a warm period in May 2013) or in controlled growth chambers (all other experiments). The growth conditions were generally set as: long days (16h light/ 8h dark), 23°C light / 20°C dark, 120-140 μmol m^-2^ s-^1^ and 65% humidity, or 16°C light / 16°C dark for ambient temperature experiment. Vernalization treatment was 3 weeks at 4°C.

### Phenotype scoring

For the genome-wide association (GWA), six randomized replicates per accession were analyzed. For each plant, the rosette diameter was measured from a pair of diameters of four biggest leaves, after six weeks of growth. For the flowering time quantification, 3 replicates were recorded for each accession. The flowering time was quantified as the growth days required for a 1 cm visible bolt and leaf number at bolting stage. 100 days were set as the flowering time for the ultimate non-flowering accessions. In mutant experiments, 5-17 plants were analyzed at bolting stage (1 cm visible bolt) for rosette diameter, leaf number and flowering days as previously described (Lempe *et al*., 2005; Salome *et al*., 2011). The Zn effect was calculated as: (plus Zn-minus Zn)/plus Zn x 100.

### Genome-wide association (GWA)

GWA was conducted using the multi-locus mixed-model to overcome the influence of population structure (Segura *et al*., 2012) with previously determined kinship matrix and SNP data from either fully sequenced genotypes or microarray analysis (Stetter *et al*., 2015). Only 162 accessions were used in GWA mapping, as the SNP data of the other 6 accessions were not available. Gene enrichment close to significant SNPs located within +/-20 kb of the significant SNPs was quantified. TAIR 10 was used as the reference database (http://www.arabidopsis.org/).

### Zinc concentration determination

6-week-old plants were harvested, dried in 60°C for a week and milled. Around 0.1 g milled materials were digested with 2.5 ml 69 % HNO_3_ and 2 ml 30% HCl for 1 hour. The samples were placed in a microwave at 170 °C for 25 minutes, followed by 200 °C for 40 minutes. The extract was measured by atomic absorption spectrometry (Thermo Fisher Scientific, United Kingdom) to determine the tissue Zn concentration.

### Quantitative RT-PCR analysis

Plants were grown in +Zn and -Zn soil-sand mixes (described above) with 3-5 replicates. For each replicate, 10-20 seedlings were harvested and pooled at around noon at 14 DAS (days after sowing, not yet bolting) with liquid nitrogen, before storing in -80°C. Total RNA of all seedlings was extracted with the innuPREP Plant RNA Kit (Analytik Jena, Germany) after plants were homogenized (Retsch, Germany). Around 1 μg total RNA was used to synthesize a cDNA library using the QuantiTect Reverse Transcription Kit (Qiagen, Germany). Gene-specific primes for qRT-PCR were designed according to the *Arabidopsis* genome sequence information TAIR10 (https://www.arabidopsis.org/) and Primer-BLAST (http://www.ncbi.nlm.nih.gov/tools/primer-blast/), quality-checked by using PCR Primer Stats (http://www.bioinformatics.org/sms2/pcr_primer_stats.html). Primers were ordered from Life Technologies (Darmstadt, Germany) and listed in the Supplemental Table S5. For the PCR procedure, 15 μl reaction mix was used, containing 6 μl 20x diluted cDNA, 7.5 μl SYBR Green Supermix (KAPA Biosystems, United States), 0.3 μl forward primers, 0.3 μl reverse primers and 0.9 μl RNase-free H_2_O. The reaction was conducted in 384-well plates in an RT-PCR system (Bio-Rad, München, Germany). The standard protocol was set as: 3 min at 95 °C, followed by 44 cycles of 3 s at 95 °C and 25 s at 60 °C, and then 5 s at 65 °C for the melt curve. For all gene expression calculations, two reference genes, *SAND* (*AT2G28390*) and *PDF2* (*AT1G13320*), were used and data were normalized to the first replicate of +Zn. These two genes did not change their expression level between different Zn treatments. Reactions were performed in 3 technical replicates and 3-5 biological replicates. Relative transcript levels were calculated with the 2-ΔΔCT method by the Bio-Rad software (Livak and Schmittgen, 2001). All kits described here were used according to the manufacturer’s instructions.

### Histological analysis

Palisade cell sizes were measured as previously described (Sicard et al., 2015). Briefly, first four leaves were harvested and fixed overnight at 4 °C in FAA solution (20 ml formalin, 10 ml acetic acid, 100 ml alcohol, and 70 ml water), and dehydrated through a series of 70, 80, 90, 100% ethonal, with 5-min incubation per step. Then the samples were transferred into acetone for 5-min incubation at 95 °C, and cleared overnight in the clearing solution (100 g chloral hydrate, 10 g glycerol, and 25 ml water). Finally the samples were stained with 10 μg ml^-1^ propidium iodide for two days, and imaged with the confocal microscope (LSM700, Carl Zeiss, Germany). Four regions were measured for every leaf and cell size was averaged from four leaves. Three biological replicates were performed.

### Statistical analysis

Data analysis, graphs and statistics were done by using Microsoft Excel and R (https://www.r-project.org/). The significant differences of means for all traits in this study were performed by t-test. Multiple comparisons were done using Tukey HSD method in R. Broad-sense heritability was calculated as genotypic variance divided by total variance (Visscher *et al*., 2008). The total variance was partitioned into genetic variance and residuals.

**Supplemental data** are given online.

## Author contributions

X.C. and U.L. conceived the experiment; X.C. performed the experimental work; X.C. and U.L. analyzed data; X.C. and U.L. wrote the paper.

## Acknowledgments

We thank Karl Schmid (Stuttgart, Germany) for all accessions seeds, Markus Schmid (Umea, Sweden) for mutant seeds, Dr Huaiyu Yang for initial help with lab work, Benjamin Neuhäuser for critical reading of the manuscript and Dominik Hedderich for help with determination of flowering times. We also thank the China Scholarship Council for support.

## Supplementary Data

**Supplementary Figure S1:** The worldwide population distribution of 168 *Arabidopsis* accessions used in this study. Every red dot represents one accession.

**Supplementary Figure S2:** Relationship between rosette diameter in +Zn, -Zn and Zn effect.

**Supplementary Figure S3:** QQ-plot of GWA mapping for Zn effect.

**Supplementary Figure S4: SNP evaluation of the GWAS for Zn sensitivity. A** and **B**, Zn sensitivity of allele adenine (A) and allele guanine (G). **C**, LD values (r^2^) between identified SNP (*1G_24327565*) and *FT* (*AT1G65480, Chr1:24331428..24333934*). Two SNPs located in the gene body of *FT* were also presented. r^2^ was 0.491 between *1G_24327565* and *1G_24333548*. r^2^ lower than 0.3 was colored with blue.

**Supplementary Figure S5:** *SOC1* transcript levels in negative-response (Col-0, Po-0, Ct-1, No-1) and positive-response accessions (Lerik1-3, Koz-2, Sf-2, Cvi-0). Data were referenced to reference genes *SAND* and *PDF2*. Values were mean + SD. * denotes p<0.05.

**Supplementary Figure S6:** Relative expression levels of typical Zn deficiency-responsive genes in Col-0 and Sf-2 leaves.

**Supplementary Table S1: List of all accessions used in this study.**

**Supplementary Table S2: Summary of rosette diameter in +Zn and -Zn.**

**Supplementary Table S3: One-way ANOVA of rosette diameter in +Zn and -Zn.**

**Supplementary Table S4: Significant SNPs identified in GWA and enriched genes located +/- 20 kb of the SNPs.**

**Supplementary Table S5: List of primers used in qRT-PCR.**

